# Characterizing the relationship between the chemical structures of drugs and their activities on primary cultures of pediatric solid tumors

**DOI:** 10.1101/2020.10.31.363218

**Authors:** Saw Simeon, Ghita Ghislat, Pedro J. Ballester

## Abstract

Better drugs are required to manage pediatric cancers. A high-throughput screen of drugs in primary cultures derived from orthotopic patient-derived xenografts (O-PDX) of pediatric solid tumours has been recently published. Here we analyzed these data sets to find out whether it is possible to leverage them for identifying new drug leads in a phenotypic manner. We found that drugs bearing a higher number of heterocyclic rings, two carbon-hetero bonds and halogens are associated to submicromolar potency in alveolar rhabdomyosarcoma and osteosarcoma O-PDXs. Furthermore, Murcko scaffolds 1-cyclopentyl-octahydro-1H-indene and tetradecahydroanthracene can be utilized as starting scaffolds to selectively optimize potency against osteosarcoma since drugs bearing this scaffold displayed superior O-PDX culture potency. Lastly, we have generated QSAR (Quantitative Structure–Activity Relationship) models able to predict the potency of drugs on each O-PDX tumor. To permit their use to guide drug repositioning on these 30 O-PDX cell cultures, we are providing a user-friendly web server implementing these QSAR models at https://rnewbie.shinyapps.io/Shobek-master

## 1. INTRODUCTION

A large number of oncology therapeutic candidates have been evaluated on cancer cell line panels. This is useful to identify which of these target-active molecules have also potent phenotypic activity (e.g. cell growth inhibition) and can therefore progress further as drug candidates. For example, the proteasome inhibitor Bortezomib approved to treat various cancer types such as multiple myeloma originates from the NCI-60 screen^1^.

Until recently ^2^, there was a dearth of available samples and models of recurrent solid tumors for pediatric cancers, which hampered the application of this phenotypic screening approach on these cancer types. More drug candidates are sought after given that 20% of children with these cancers do not survive ^3^ and current drug treatments can cause long-term toxicity in the remaining 80% of children ^4^. These authors ^2^ generated pediatric cancer cell lines for several tumor types.

Another contribution of that study ^2^ is providing more advanced *in vitro* models than immortalized cell lines. Indeed, cell lines have significant limitations in their ability to model the biology and therapeutic responsiveness due to multiple reasons, e.g. DNA and mRNA of cell lines continuously change in culture, cell lines do not longer retain the intra-tumor heterogeneity present in the primary cancer, or do not contain the relevant components of the tumor microenvironment ^5^. To circumvent these limitations, pediatric cancer Orthotopic Patient-Derived Xenografts (O-PDXs) were also established within immunocompromised mice ^2^. O-PDX-derived primary cell cultures are a promising preclinical model to better predict drug response of patients in drug development and in recurrent cancer Thus, drug screening on O-PDX primary cultures constitutes a more predictive in vitro model of patient response to drugs than if made on cancer cell lines.

Overall, these authors ^2^ provided one of the most comprehensively profiled repositories to date in drug screening for pediatrics solid tumors. More concretely, 158 drugs were tested *in vitro* on 53 pediatric cancer models (23 cell lines and 30 O-PDX primary cultures). This dataset can be reused to identify which chemical scaffolds are associated with more selective, promiscuous or active drugs with views to exploiting this information for screening library design. Furthermore, *in silico* models built with Machine Learning (ML) from this dataset can be investigated to find out whether it is possible to predict drug response on these new 30 O-PDX pediatric cancer models.

Given the availability of molecules profiles for these cancer models, ML models could be generated to predict drug response on additional forthcoming patient tumors profiled in the same way. This has been investigated on cancer cell lines ^6,7^ or directly patients ^8,9^. Here we focus instead on predicting the activities of unseen molecules on each of the 30 cancer models. In particular, we will build and evaluate Quantitative structure activity relationship (QSAR) models on these datasets, which do not require any tumor profile to be determined.

QSAR is a widely applied computational method to hit identification based on the notion that structurally similar compounds exert similar activities on their biomolecular targets. QSAR may considerably reduce cost of synthesizing compound, avoid animal testing and speed up managerial decision ^10^. Generally, chemical structures are described as a set of numerical descriptors or features, which then correlate with bioactivity to establish QSAR model to predict bioactivities of novel compounds. In cancer drug discovery, QSAR has been successfully utilized in the discovery of novel inhibitors of histone deacetylase 1 (HDAC1) in which the inhibitor activity was confirmed experimentally ^11^. Similarly, novel inhibitors for mammalian target of rapamycin (mTOR) were discovered using optimal QSAR models ^12^.

Larger training datasets typically results in more predictive machine learning models. However, this does not mean that models trained on moderately-sized datasets cannot be predictive. For example, Kumar et al. successfully employed drugs and built QSAR models for 16 pancreatic cancer cell lines, with that for the YAPC and PSN1 cell lines obtaining the largest and smallest R^2^ values (0.79 and 0.42, respectively). The average number of training drugs across each cell lines was 97, indicating that these amounts of drugs in the dataset are sufficient to build QSAR models ^13^. Here we will be able to employ 50% more training drugs per tumor.

To sum up, in an attempt to understand the underlying relationship between drug responses in O-PDXs and molecular structure, we have analyzed a recent flood of pediatric cancer data ^2^, specifically on its O-PDXs repository for pediatrics solid tumors. As such, it represents an ideal open-access database to perform cheminformatics analyses. We have used these data to investigate the relationship between: first, structure space covered by drugs for pediatric cancer; second, O-PDXs potency and chemical features; third, selectivity and promiscuous drugs for pediatric cancers.

## 2. MATERIALS AND METHODS

### 2.1. O-PDXs Data

The bioactivity of a drug on a tumor model was measured as the effective molar concentration of the drug required to achieve a 50% of tumour growth decrease (EC_50_). EC_50_ values are converted to pEC_50_ values via the base10-logarithmic transformation (drug-model pairs with pEC_50_ < 2 were not considered given their low reliability). The employed datasets come from screening drugs on pediatric cancer O-PDX-derived primary cell culture and cell lines drug sensitivity data from Stewart et al. ^2^. As shown in supplementary table S1 containing high-throughput drug screening data,

The 30 cell cultures derived from O-PDXs can be classified according to tumor type: rhabdoid tumor (RT), embryonal rhabdomyosarcoma (RMS_E), alveolar shabdomyosarcoma (RMS_A), osteosarcoma (OS), neuroblastoma (NB) and Ewing sarcoma (EWS). Characteristics of drug response in each cell cultures derived from the 30 O-PDXs are shown in Table S2 in which minimum and maximum pEC_50_ of a drug are 2.55 and 10.00, respectively.

The pEC_50_ values of each tumor type are grouped into classes (Active, Intermediate and Inactive). Drugs displaying pEC_50_ above 6 (i.e. EC_50_ values below 1 µM) are classified as actives whereas drugs exerting values below 5 (corresponding to EC_50_ values greater than 10 µM) were classified into inactive and those with intermediate potency with pEC_50_, ranging from 5 to 6, were not considered for the chemical scaffold analysis because of their dubious nature. The ratio of inactives against actives (Figure 2) is employed to identify whether a O-PDX primary cultures would need focus on the development of additional drugs (a value < 1, indicates there are more active drugs than inactive ones and vice versa for values > 1). O-PDX primary cultures model where categorized based on the ratio values of >1 and <1 in which 18 O-PDXs have <1 while the remaining 12 have ratio of >1.

**Figure 1:**
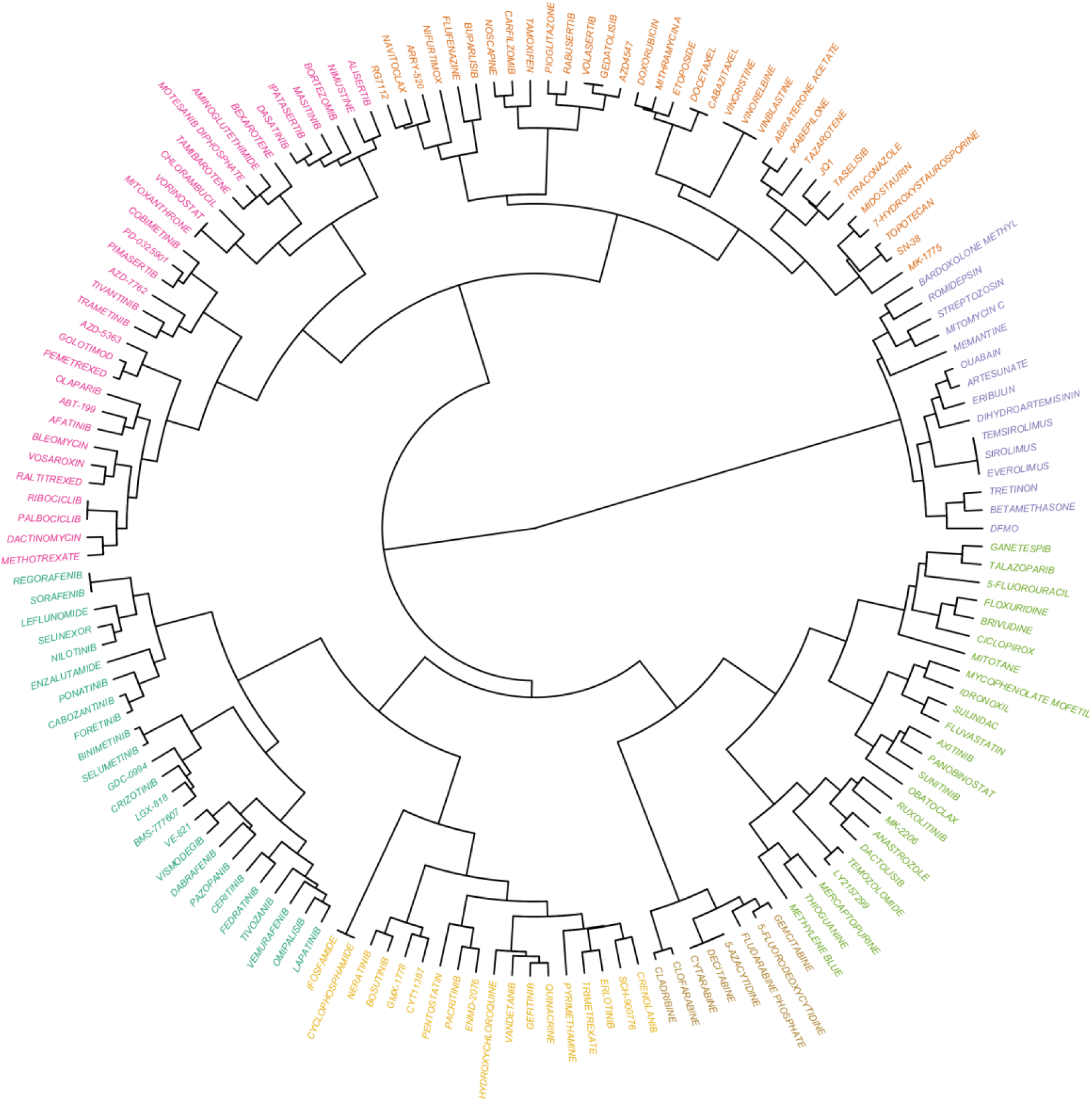
Hierarchical clustering of the tested drugs using E-state fingerprints showing seven clusters. Each cluster colored differently.

**Figure 2:**
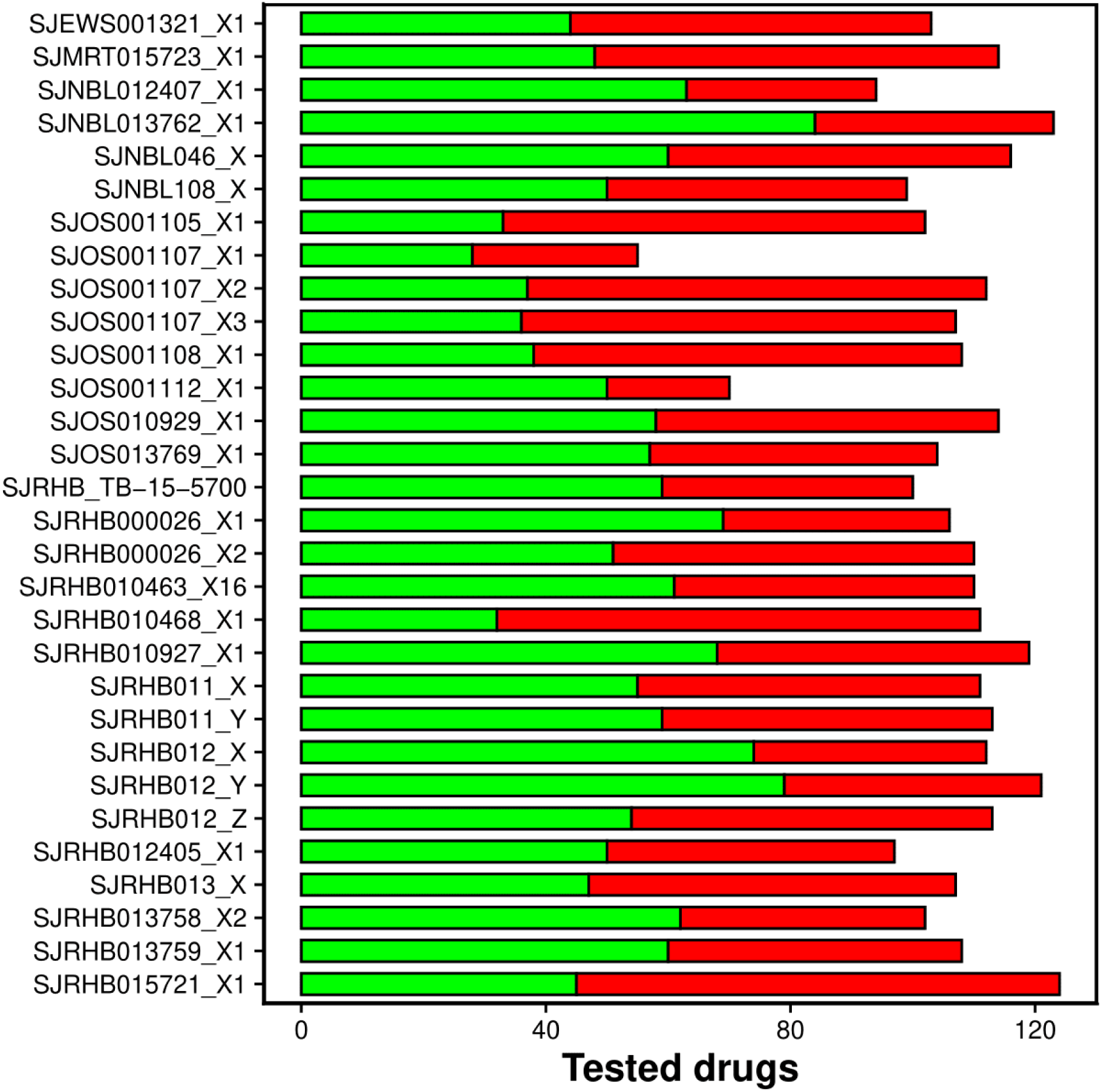
Number of actives (green) and inactives (red) among the tested drugs per PDOX-derived cell line.

### 2.2. Feature generation

Drugs are treated with chemical curation workflow with the following steps: (i) removal of inorganic and mixture, (ii) structural conversion and cleaning, (iii) normalization of specific chemotype, (iv) removal of duplicate and (v) final manual checking ^14^. The ChemAxon Standardizer was utilized to treat drugs with the following options: Strip Salt, Aromatize, Clean 3D, Tautomerize, Neutralize and Remove explicit hydrogen ^15^. After this curation process, 155 of the 158 drugs were retained.

Two chemical structure fingerprints for each drug were calculated. The first bit-based fingerprint using the PaDEL-Descriptor software ^16^ and showing the presence or absence of a various substructures (307 features per drug). The E-state fingerprint was also employed, providing counts of 79 E-state atom types per drug associated with the Kier-Hall electronegativity, graph topology and functional group (calculated using the rcdk R package), which encode the Electrotopological state. Lastly, 6 physico-chemical descriptors are calculated using JChem Base: lipophilicity (logP), geometric counts (polar surface area or PSA, chiral center count), H-bond donor (HBD) and acceptor counts (HBA) and molecular weight (MW). After removing near-zero-variance features and intercorrelations of features higher than 7 (using the nearZero and findCorrelation functions of the R caret package), 40 features per drug were retained.

From the preliminary clustering analysis at a cutoff of Tanimoto similarity 0.85, 134 singletons were obtained, indicating that the chemical space of these drugs is highly diverse. Visual inspection shows that there are 7 major clusters for a broad range of cutoffs. Figure 1 showed a hierarchical clustering dendrogram with those 7 clusters comprising the 155 drugs.

### 2.3. Data Splitting

Each O-PDX cell culture data set was partitioned into training (80%) and test (20%) sets using the Kennard-Stone algorithm ^17^ to prevent unforeseen biased selection. Briefly, the method selects a subset of instances providing a uniform coverage of the feature space to act as a test set. Test set building begins by finding two instances, which are farthest away using their Euclidean distance in feature space. The remaining instances are sequentially added to the test set by selecting that with the highest distance to any of the test instances so far. This process is repeated until the required numbers of instances for the test set is reached. The non-selected instances are employed as a non-overlapping training set.

### 2.4. Predictive modelling

Random forest (RF) is an ensemble method in which several decision trees (DT) are used to produce a single consolidated model that is more powerful than the individual constituent DT ^18^. RF used different samples of the same data set to train multiple versions of the model. These models then vote on the correct answer for a new observation and an average decision is made, which is known as bagging or bootstrap aggregation. Thus, it combines the results from different models that are built on the same data set, by using majority voting. RF creates a random sample size n, in which n is the number of observations (i.e. cell-tested drugs) in the original data set with replacement meaning that some of the observations from the original training set will be repeated and some will not be chosen at all, also known as bootstrap sampling. From the original data set, observations that were not chosen for a particular iteration of the procedures is known as out-of-bag (OOB) observation. These observations are not used in the training model, thus, instead of relying on the observations used to train the model at every stop, we can actually use the OOB observations to record the accuracy of a particular model. Then the average over all the OOB accuracy rates can be used to obtain an average accuracy. This average accuracy is more likely to be a realistic and an objective estimate of the performance of the bagged classifier on unseen data, for a particular observation, the dependent variables are thus decided as the majority vote over all individual models for which the observation was not picked to their training samples. The samples generalized from sampling the original data set with replacement, known as bootstrapped samples, are similar to drawing multiple samples from the same distribution. By trying to estimate the same target function using a number of different samples instead of just one, the averaging process reduces the variance of the result as well as the model. RF models have been used extensively to predict protein-ligand binding affinities on macromolecular targets^19–22^ or the activity/synergy of molecules on cell lines ^23–26^.

For comparison, we also applied Partial Least Squares (PLS). PLS is a linear regression model which seeks to relate observations to predict the pEC_50_ based on the characteristics of variables. The idea behind PLS is to produce a small number of linear combinations of the original variables and use these linear combinations as input to perform a regression model. The latent variables can be assumed to account for most of the variance in the dataset in which descriptors are highly correlated. Therefore, it seeks latent variabels that have high variance and high correlation with the dependent variable by simultaneously projecting the descriptors into latent component to correlate with bioactivity values. PLS models have been commonly used to predict the ADMET properties ^27^ and in vitro potency of protein-ligand binding affinities ^28^.

### 2.5. Assessment parameters

Model validation is an important process in empirical modeling. There are several statistical assessment tools that can be used to assess the effectiveness and efficiency. In this study, the square of correlation coefficient (R^2^), cross-validation coefficient and external validation. The R^2^ shows the percent variation of the model in which the value closer to 1 represents the better predictive model. The R^2^,Q^2^_cv_ and Q^2^_test_ can be calculated using the following equations:

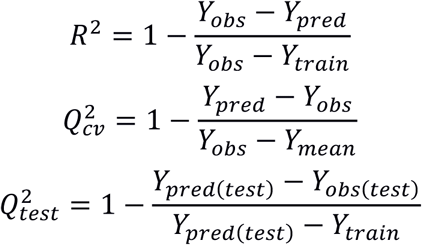

in which Y denotes the dependent variable, which is drug-induced growth inhibition on a particular tumor culture or cell line. Leave-One-Out Cross-Validation (LOOCV) and external testing was used to validate reliability of the predictive models. As usual, LOOCV was utilized to estimate the generalization ability of the model using the training set, with the external test set providing further assessment of the extrapolation power of the model. LOOCV was performed on the training where the data are separated into different observations. One drug is left out as test fold, where the rest are used for training models. This process is repeated iteratively until all the data samples are in the test set exactly once. External validation evaluates the robustness of the predictive model by serving as unseen data not used to build the model.

### 2.5. Molecular cluster analysis

Privileged substructures or scaffolds ^29^ are structures likely to provide useful ligands for more than one target. To identify privileged scaffolds for each tumor model, we followed this protocol: 1) Each drug is represented by its Murcko scaffold ^30^ which represents its core chemical structure; 2) Murcko scaffolds contained by at least 3 drugs were retained for further analysis; 3) mean pEC_50_ of these scaffolds were compared to that of drugs tested on the tumor type; 4) Murcko scaffolds are prioritized (‘privileged’) in terms of how large is their potency (mean pEC_50_) compared to the background of all tested drugs. Maximizing Selectivity Index (SI) implies maximizing phenotypic potency while reducing overall potency of drugs and eventually leads to reduce the risk of attrition. The combination of SI together with chemical scaffolds relevant to phenotypic potency has the potential to accelerate lead optimization.

## 3. RESULTS

### 3.1. Univariate analysis of O-PDX primary cell cultures

We started by identifying which chemical structure features discriminate between active and inactive drugs on each of the 30 tumor models. The normality of each data set was assessed using the Shapiro-Wilk test, a hypothesis that was rejected for each feature and data set at the 0.05 confidence level. Thus, the non-parametric Mann-Whitney U test was utilized to identify discriminative features for the two drug classes. Figure 2 shows that SJOS001105_X1 (OS), SJOS001107_X2 (OS) and SJRHB010468_X1 (RMS_A) have two times more inactive drugs than active ones, suggesting that more drugs active on these tumor types would be beneficial. We hence employed these three tumor models as case studies. Tables S3 to S5 show the mean and standard deviation of the chemical features on each drug class along with their *P* values (Mann-Whitney U test). Of the 40 employed features, 27.5%, 32.5% and 22.5% lead to significant *P* values for SJOS001105_X1, SJOS001107_X2 and SJRHB010468, respectively. For example, SubFC1 is the primary carbon of chemical compounds. This analysis shows that active drugs (i.e. those with potency better than micromolar) contain this substructure more often than inactive drugs (2.649 ± 2.946 vs. 0.918 ± 1.233) for SJOS001107_X2 (OS) PDOX.

### 3.2. Identification of privileged scaffolds with desirable potency in O-PDX primary cultures

The mean potency (mean pEC_50_) of rhabdoid tumor, embryonal rhabdomyosarcoma, alveolar rhabdomyosarcoma, osteosarcoma, neuroblastoma and ewing sarcoma were 5.42, 5.81, 5.73, 5.41, 5.85 and 5.39, respectively. These findings suggest that PDX drugs seldomly have nanomolar potency. The mean pEC_50_ of all test drugs were compared against mean pEC_50_ of Murcko scaffold and it was observed that the scaffold 1-cyclopentyl-octahydro-1H-indene is associated with submicromolar potency (mean pEC_50_ = 6.80 vs 5.42) for rhabdoid tumor whereas lower/comparable mean pEC_50_ were found for neuroblastoma (5.56 vs. 5.85), osteosarcoma (5.42 vs. 5.41), embryonal rhabdomyosarcoma (5.38 vs 5.81) and alveolar rhabdomyosarcoma (5.40 vs 5.73), suggesting that this scaffold is both biasedly potent and selective toward rhabdoid tumor. On the other hand, the scaffold octahydro-1H-indene is associated to submicromolar potency for rhabdoid tumor (6.14 vs 5.42), embryonal rhabdomyosarcoma (6.16 vs 5.81), alveolar rhabdomyosarcoma (6.17 vs 5.73) suggesting that it can be used as a starting point to design promiscuous drugs against multiple O-PDX primary cultures as it may be more resilient against drug-resistant mutations. A few examples can be purchased for each scaffold for future screening in designing selective drugs candidates against a particular tumour type or promiscuous drugs that display O-PDX primary culture potency across several tumour types (Figure 5), which may be useful for compound library design.

### 3.3. Predicting the potencies of drugs on O-PDX primary cultures

QSAR were generated with simple physicochemical descriptors and fingerprint descriptors using RF. The performance of this algorithm in LOOCV and external test set can be visualized across the 30 O-PDX models in Figures 3 and 4, respectively. The results for the 30 O-PDX primary cultures are fully reported in Table S6 in which the corresponding prediction error is given from the RMSE ranges from 0.38 to 0.70. The cross-validated RMSE ranges from 0.62 to 1.18, indicating that the models are robust. The model performs equally well on the test set for all O-PDX primary cultures which is unsurprising given it was partitioned using Kennard-Stones. The Rs (Spearman correlation) of the test set ranges from 0.82 to 0.99. In an effort to put RF model performance in perspective, the results of the PLS models were compared. The results of model generated using PLS are reported in Table S7. The results suggest that RF is more effective than the PLS on account of large R^2^ and lower RMSE values in fit. The R^2^ ranges from 0.18 to 0.76 for the training set. All RF models display lower prediction error (i.e., RMSE) than those found for PLS, consistent with the previous reports that RF tend to perform better than linear models^31^.

**Figure 3:**
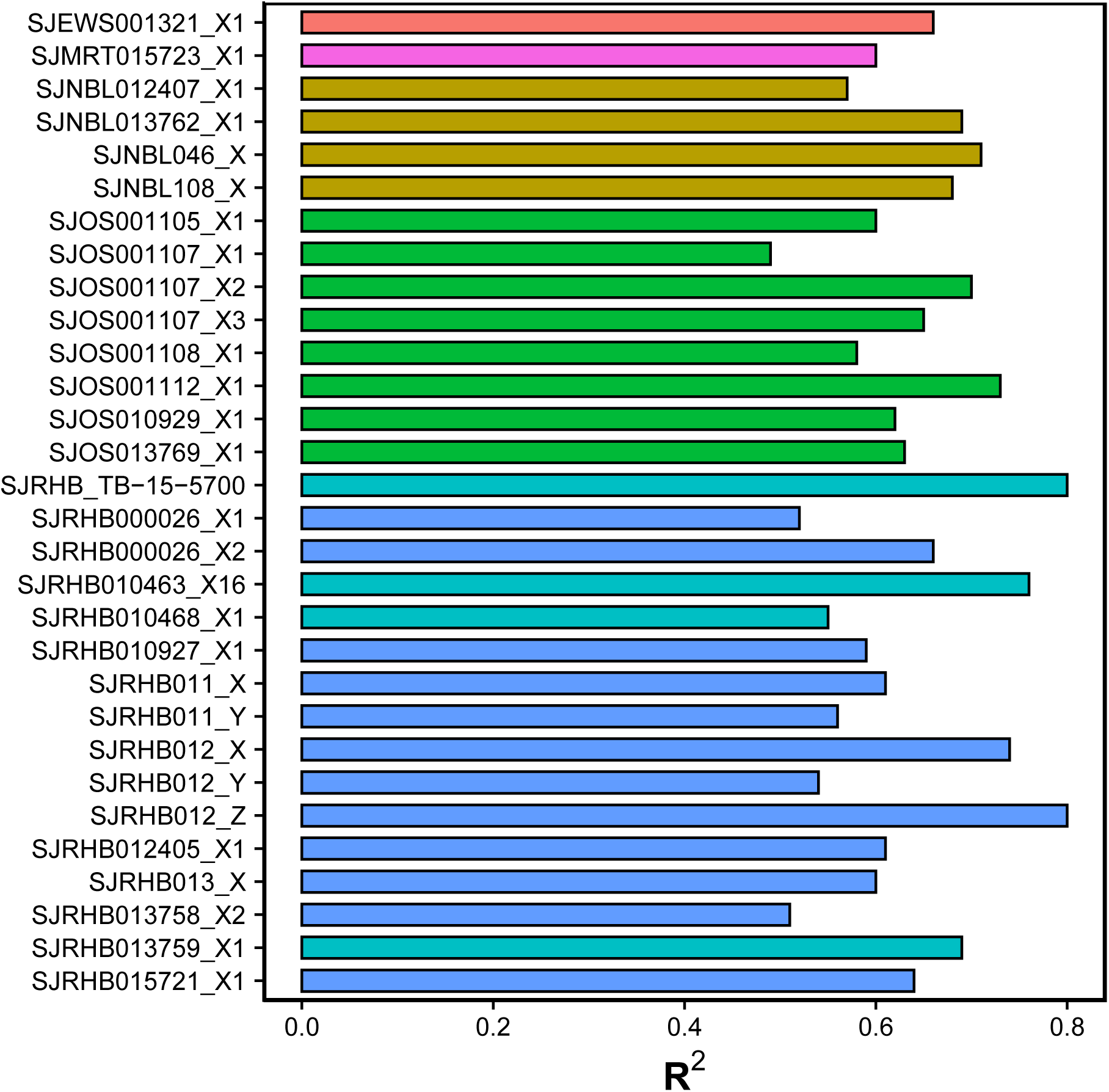
Predictive performance of the Random Forest model for each PDOX cell line as given by the R^2^ of the leave-one-out cross-validation. The colors represent the different pediatric cancer types.

**Figure 4:**
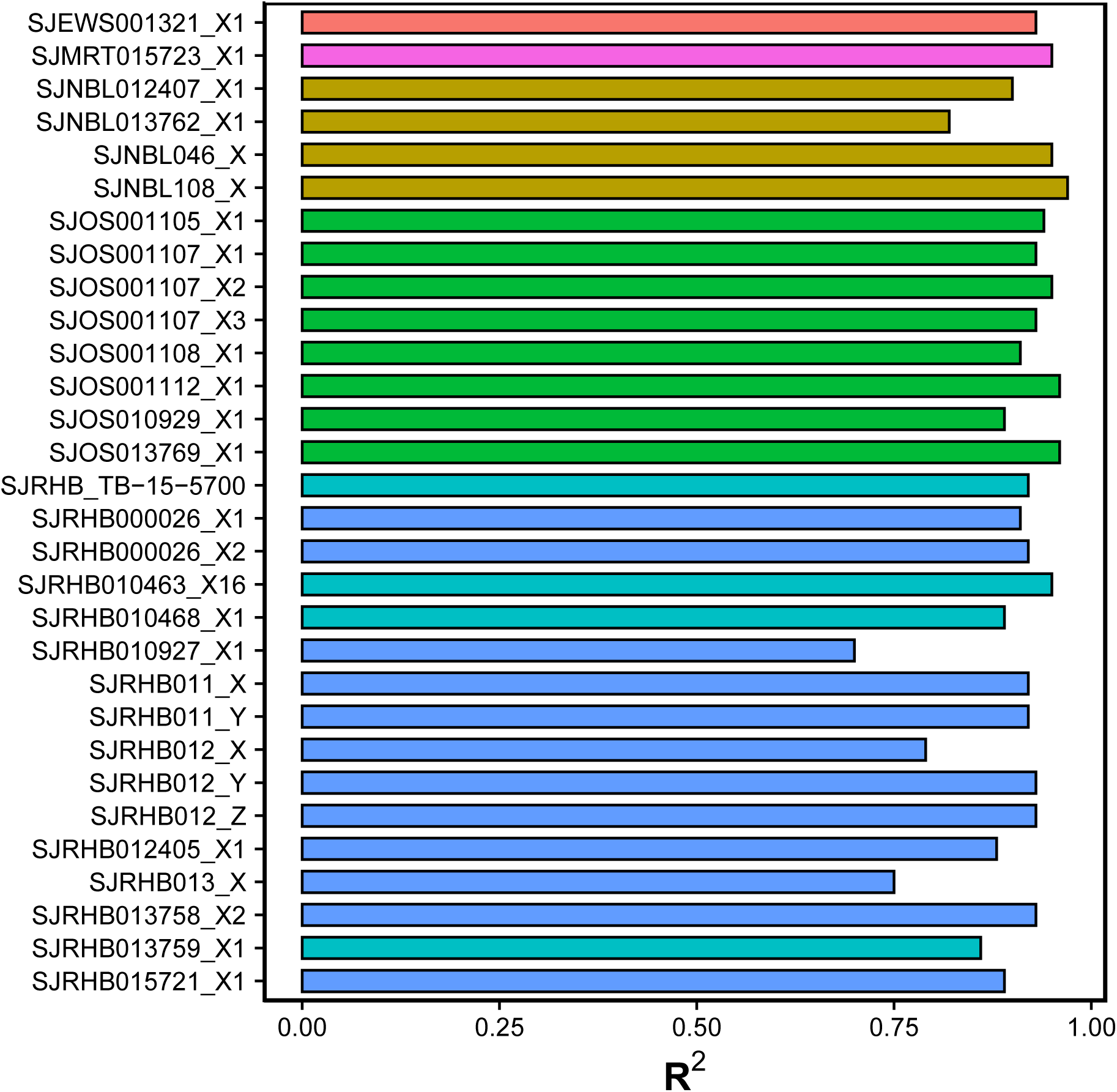
Predictive performance of the Random Forest model for each PDOX cell line as given by the R2 on the test set. The colors represent the different pediatric cancer types.

**Figure 5:**
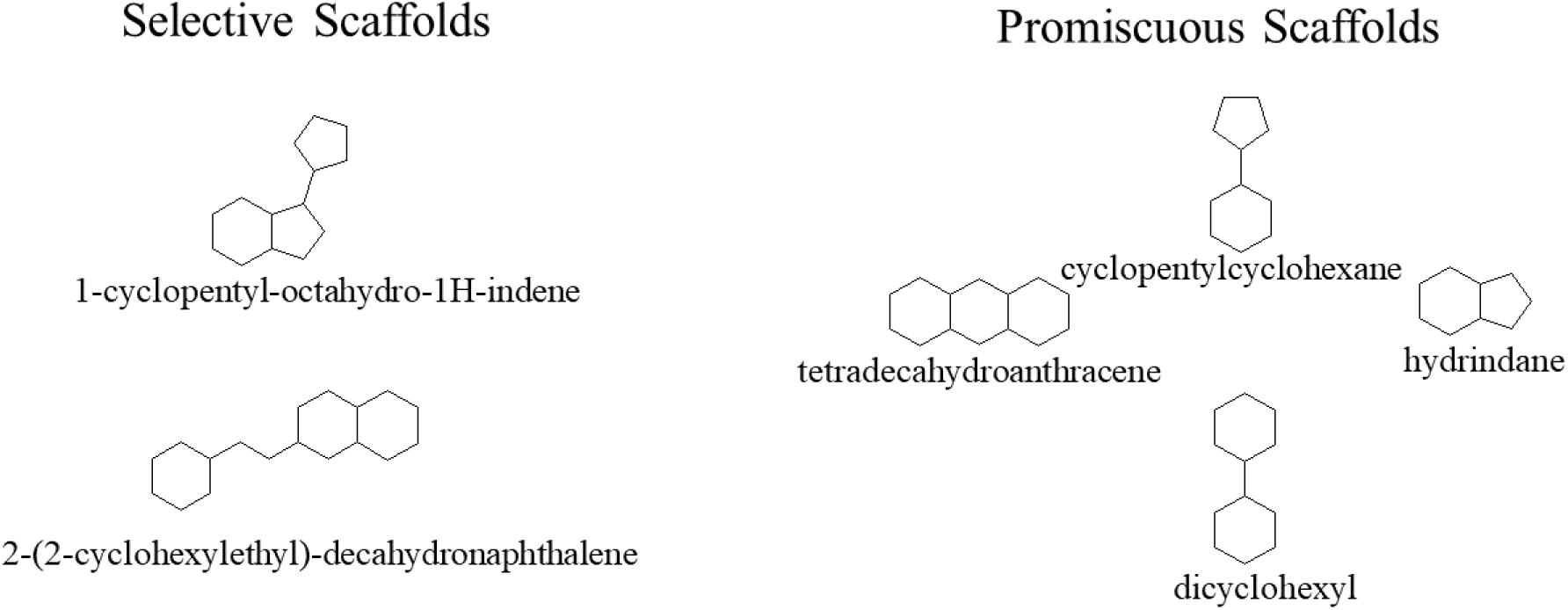
Structure of selected scaffolds for future screening in designing selective or promiscuous drugs candidates. The IUPAC names were obtained using structure to name tool from ChemAxon MarvinSketch (Product version: 20.19.0)

## CONCLUSION

There is an urgent need for efficient and effective tools to guide the identification of new drug leads to develop better cancer therapies. O-PDX-derived primary cell cultures are highly-valuable preclinical model with which to identify these leads. We have identified substructures and murcko scaffolds with potent selective activity towards specific O-PDX primary cultures as well as those with promiscuous activity across the 30 O-PDX cultures. This can be used as a guide in the early drug discovery process to detect whether a chemical series is likely to achieve the necessary potency for these tumor models. Furthermore, QSAR has been successfully applied in hit identification and lead optimization. Here we have trained and validated RF-based QSAR models providing excellent performance across O-PDX primary cell culture models. To facilitate the repositioning of drugs on any O-PDXs, we provide all 30 predictive models at https://rnewbie.shinyapps.io/Shobek-master.

## Supporting information

All tables

## AUTHOR CONTRIBUTIONS

P.J.B. designed and wrote the paper with the assistance of S.S. and G.G. S.S. performed experiments and prepared figures. All authors commented on the article and approved the final submitted article.

## ACKNOWLEDGEMENTS

Standardizer was used for structure canonicalization and transformation, JChem Version: 20.17.0, 2020, ChemAxon (http://www.chemaxon.com).

