## Supplementary material for "Characterizing the relationship between the chemical structures of drugs and their activities on primary cultures of pediatric solid tumors": All tables

**ARTICLE TYPE**

Table S1: Characteristics of the drug-model bioactivity matrix

| Characteristics | PDOX primary cultures |
|---|---|
| $EC_{50}$ values | 4626 |
| Drugs (observations) | 158 |
| Unique tumor models | 30 |
| Matrix Completeness | 97.59% |

Table S2: Overview of the drugs sensitivity data for each PDOX primary culture. N is the number of tested drugs. EC50 values are in molar concentration units.

| ID | PDOX primary culture | N | $pEC_{50}$ (Mean±SD) | $pEC_{50}$ (Min) | $pEC_{50}$ (Max) |
|---|---|---|---|---|---|
| 1 | SJEWS001321_X1 | 156 | 5.61 ± 1.05 | 4.00 | 9.59 |
| 2 | SJMRT015723_X1 | 153 | 5.71 ± 1.33 | 4.00 | 10.00 |
| 3 | SJNBL012407_X1 | 139 | 6.17 ± 1.45 | 4.00 | 10.00 |
| 4 | SJNBL013762_X1 | 151 | 6.29 ± 1.48 | 4.00 | 10.00 |
| 5 | SJNBL046_X | 156 | 5.92 ± 1.39 | 4.00 | 10.00 |
| 6 | SJNBL108_X | 153 | 5.78 ± 1.31 | 4.00 | 10.00 |
| 7 | SJOS001105_X1 | 156 | 5.35 ± 1.00 | 4.00 | 10.00 |
| 8 | SJOS001107_X1 | 106 | 5.78 ± 1.39 | 4.00 | 8.87 |
| 9 | SJOS001107_X2 | 154 | 5.43 ± 1.11 | 4.00 | 10.00 |
| 10 | SJOS001107_X3 | 155 | 5.43 ± 1.13 | 4.00 | 10.00 |
| 11 | SJOS001108_X1 | 153 | 5.50 ± 1.29 | 4.00 | 9.12 |
| 12 | SJOS001112_X1 | 121 | 6.04 ± 1.32 | 4.00 | 10.00 |
| 13 | SJOS010929_X1 | 156 | 5.97 ± 1.52 | 4.00 | 10.00 |
| 14 | SJOS013769_X1 | 153 | 5.83 ± 1.26 | 4.00 | 10.00 |
| 15 | SJRHB_TB-15-5700 | 131 | 6.14 ± 1.69 | 4.00 | 10.00 |
| 16 | SJRHB000026_X1 | 146 | 6.29 ± 1.61 | 4.00 | 10.00 |
| 17 | SJRHB000026_X2 | 147 | 5.71 ± 1.34 | 4.00 | 10.00 |
| 18 | SJRHB010463_X16 | 158 | 5.91 ± 1.37 | 4.00 | 10.00 |
| 19 | SJRHB010468_X1 | 155 | 5.36 ± 1.04 | 4.00 | 8.66 |
| 20 | SJRHB010927_X1 | 154 | 5.99 ± 1.52 | 4.00 | 10.00 |
| 21 | SJRHB011_X | 152 | 5.86 ± 1.34 | 4.00 | 10.00 |
| 22 | SJRHB011_Y | 139 | 6.03 ± 1.87 | 2.55 | 10.00 |
| 23 | SJRHB012_X | 154 | 6.15 ± 1.46 | 4.00 | 10.00 |
| 24 | SJRHB012_Y | 147 | 6.35 ± 1.64 | 4.00 | 10.00 |
| 25 | SJRHB012_Z | 151 | 5.76 ± 1.52 | 4.00 | 10.00 |
| 26 | SJRHB012405_X1 | 147 | 5.81 ± 1.22 | 4.00 | 10.00 |
| 27 | SJRHB013_X | 152 | 5.74 ± 1.34 | 4.00 | 10.00 |
| 28 | SJRHB013758_X2 | 146 | 6.10 ± 1.49 | 4.00 | 10.00 |
| 29 | SJRHB013759_X1 | 150 | 5.82 ± 1.28 | 4.00 | 9.75 |
| 30 | SJRHB015721_X1 | 158 | 5.52 ± 1.35 | 4.00 | 10.00 |

Table S3: Summary of the mean and standard deviations of substructure fingerprints along with their p-values derived from the statistical difference test using the Mann-Whitney U test for drugs tested on an OS PDOX primary culture (SJOS00105_X1)

| Features | Description | Active drugs | Inactive drugs | p-value |
|---|---|---|---|---|
| nAromAtom | Number of aromatic atoms | 7.031 ± 8.197 | 10.899 ± 8.845 | 0.028 |
| SubFPC52 | Imine | 0.125 ± 0.336 | 0.000 ± 0.000 | 0.003 |
| SubFPC127 | Peptide middle | 0.250 ± 1.107 | 0.000 ± 0.000 | 0.038 |
| SubFPC135 | Carboxyl derivative | 0.844 ± 1.247 | 0.275 ± 0.591 | 0.012 |
| SubFPC136 | Vinylogous acid | 0.156 ± 0.515 | 0.000 ± 0.000 | 0.011 |
| SubFPC169 | Phenol | 0.250 ± 0.672 | 0.014 ± 0.120 | 0.016 |
| SubFPC172 | Arylfluoride | 0.063 ± 0.246 | 0.377 ± 0.729 | 0.018 |
| SubFPC181 | Hetero N nonbasic | 0.688 ± 0.931 | 1.420 ± 1.344 | 0.009 |
| SubFPC188 | Nitro | 0.063 ± 0.246 | 0.000 ± 0.00 | 0.038 |
| SubFPC297 | Anion | 0.063 ± 0.246 | 0.000 ± 0.00 | 0.038 |
| SubFPC299 | Salt | 0.063 ± 0.246 | 0.000 ± 0.00 | 0.038 |

Table S4: Summary of the mean and standard deviations of substructure fingerprints along with their p-values derived from the statistical difference test using the Mann-Whitney U test for drugs tested on an OS PDOX primary culture (SJOS001107_X2)

| Features | Description | Active drugs | Inactive drugs | p-value |
|---|---|---|---|---|
| SubFPC1 | Primary carbon | 2.649 ± 2.946 | 0.918 ± 1.233 | 0.001 |
| SubFPC3 | Tertiary carbon | 0.7575 ± 1.188 | 0.178 ± 0.586 | 0.002 |
| SubFPC4 | Quaternary carbon | 0.541 ± 1.169 | 0.192 ± 0.593 | 0.047 |
| SubFPC5 | Alkene | 0.568 ± 1.015 | 0.192 ± 0.430 | 0.026 |
| SubFPC23 | Amine | 0.108 ± 0.393 | 0.425 ± 0.780 | 0.014 |
| SubFPC26 | Tertiary aliphatic amine | 0.081 ± 0.277 | 0.315 ± 0.621 | 0.044 |
| SubFPC49 | Ketone | 0.432 ± 0.801 | 0.055 ± 0.283 | 0.000 |
| SubFPC52 | Imine | 0.108 ± 0.315 | 0.000 ± 0.000 | 0.005 |
| SubFPC100 | Secondary amide | 0.243 ± 0.796 | 0.384 ± 0.738 | 0.035 |
| SubFPC127 | Peptide middle | 0.270 ± 1.071 | 0.000 ± 0.000 | 0.015 |
| SubFPC276 | Epoxide | 0.054 ± 0.229 | 0.000 ± 0.000 | 0.047 |
| SubFPC287 | Conjugated double bond | 3.162 ± 2.421 | 2.219 ± 2.422 | 0.026 |
| SubFPC303 | Michael acceptor | 0.541 ± 1.043 | 0.233 ± 0.657 | 0.037 |

Table S5: Summary of the mean and standard deviations of substructure fingerprints along with their p-values derived from the statistical difference test using the Mann-Whitney U test for drugs tested on a RMS_A PDOX primary culture (SJRHB010468_X1)

| Features | Description | Active drugs | Inactive drugs | p-value |
|---|---|---|---|---|
| nRing | Number of Ring | 4.677 ± 1.536 | 3.654 ± 1.704 | 0.001 |
| SubFPC52 | Imine | 0.097 ± 0.301 | 0.013 ± 0.113 | 0.037 |
| SubFPC86 | Lactone | 0.161 ± 0.454 | 0.026 ± 0.159 | 0.033 |
| SubFPC99 | Primary amide | 0.065 ± 0.249 | 0.000 ± 0.000 | 0.025 |
| SubFPC169 | Phenol | 0.161 ± 0.523 | 0.000 ± 0.000 | 0.006 |
| SubFPC180 | Hetero N basic no H | 0.484 ± 0.769 | 0.103 ± 0.305 | 0.001 |
| SubFPC184 | Heteroaromatic | 2.258 ± 1.731 | 1.513 ± 1.448 | 0.035 |
| SubFPC275 | Heterocyclic | 4.194 ± 2.372 | 3.026 ± 1.758 | 0.018 |
| SubFPC307 | Chiral center specified | 22.129 ± 7.504 | 17.962 ± 7.076 | 0.009 |

Table S6: Summary of the predictive performance (Random Forest) on each PDOXs cell line for training set, Leave-One-Out Cross-Validation (LOOCV) and test set.

| PDOXs cell line | Training Set | | | LOOCV | | | Test set | | |
|---|---|---|---|---|---|---|---|---|---|
| | R (Spearman) | $R^2$ | RMSE | R (Spearman) | $R^2$ | RMSE | R (Spearman) | $R^2$ | RMSE |
| SJEWS001321_X1 | 0.95 | 0.91 | 0.42 | 0.77 | 0.63 | 0.67 | 0.95 | 0.93 | 0.54 |
| SJMRT015723_X1 | 0.93 | 0.90 | 0.52 | 0.75 | 0.58 | 0.86 | 0.84 | 0.88 | 0.61 |
| SJNBL012407_X1 | 0.94 | 0.90 | 0.57 | 0.81 | 0.60 | 0.89 | 0.97 | 0.95 | 0.63 |
| SJNBL013762_X1 | 0.96 | 0.92 | 0.57 | 0.72 | 0.56 | 0.93 | 0.93 | 0.90 | 0.62 |
| SJNBL046_X | 0.91 | 0.87 | 0.60 | 0.80 | 0.70 | 0.78 | 0.90 | 0.82 | 0.68 |
| SJNBL108_X | 0.95 | 0.92 | 0.47 | 0.82 | 0.71 | 0.71 | 0.93 | 0.95 | 0.60 |
| SJOS001105_X1 | 0.94 | 0.92 | 0.38 | 0.77 | 0.66 | 0.62 | 0.95 | 0.97 | 0.59 |
| SJOS001107_X1 | 0.91 | 0.89 | 0.57 | 0.79 | 0.69 | 0.80 | 0.92 | 0.86 | 0.56 |
| SJOS001107_X2 | 0.92 | 0.89 | 0.44 | 0.60 | 0.60 | 0.70 | 0.90 | 0.94 | 0.66 |
| SJOS001107_X3 | 0.94 | 0.91 | 0.44 | 0.68 | 0.47 | 0.81 | 0.92 | 0.93 | 0.49 |
| SJOS001108_X1 | 0.89 | 0.89 | 0.56 | 0.71 | 0.69 | 0.81 | 0.94 | 0.95 | 0.82 |
| SJOS001112_X1 | 0.95 | 0.92 | 0.47 | 0.82 | 0.64 | 0.78 | 0.91 | 0.89 | 0.59 |
| SJOS010929_X1 | 0.94 | 0.92 | 0.61 | 0.80 | 0.65 | 0.92 | 0.95 | 0.93 | 0.82 |
| SJOS013769_X1 | 0.95 | 0.91 | 0.49 | 0.67 | 0.57 | 0.89 | 0.90 | 0.91 | 0.56 |
| SJRHB_TB-15-5700 | 0.98 | 0.94 | 0.54 | 0.86 | 0.80 | 0.83 | 0.91 | 0.92 | 0.62 |
| SJRHB000026_X1 | 0.95 | 0.91 | 0.64 | 0.73 | 0.50 | 1.07 | 0.95 | 0.91 | 0.71 |
| SJRHB000026_X2 | 0.97 | 0.93 | 0.47 | 0.82 | 0.64 | 0.80 | 0.95 | 0.92 | 0.53 |
| SJRHB010463_X16 | 0.96 | 0.93 | 0.48 | 0.82 | 0.72 | 0.74 | 0.98 | 0.96 | 0.63 |
| SJRHB010468_X1 | 0.87 | 0.91 | 0.42 | 0.74 | 0.62 | 0.72 | 0.87 | 0.89 | 0.69 |
| SJRHB010927_X1 | 0.96 | 0.93 | 0.55 | 0.85 | 0.78 | 0.76 | 0.99 | 0.95 | 0.82 |
| SJRHB011_X | 0.92 | 0.90 | 0.56 | 0.74 | 0.54 | 0.82 | 0.95 | 0.89 | 0.44 |
| SJRHB011_Y | 0.95 | 0.92 | 0.79 | 0.75 | 0.58 | 1.18 | 0.84 | 0.75 | 0.84 |
| SJRHB012_X | 0.95 | 0.88 | 0.65 | 0.74 | 0.60 | 0.91 | 0.96 | 0.92 | 0.92 |

| | | | | | | | | | |
|---|---|---|---|---|---|---|---|---|---|
| SJRHB012_Y | 0.96 | 0.92 | 0.58 | 0.76 | 0.55 | 1.03 | 0.93 | 0.92 | 0.65 |
| SJRHB012_Z | 0.93 | 0.90 | 0.60 | 0.69 | 0.51 | 1.11 | 0.90 | 0.93 | 0.51 |
| SJRHB012405_X1 | 0.97 | 0.91 | 0.41 | 0.77 | 0.58 | 0.77 | 0.82 | 0.70 | 0.88 |
| SJRHB013_X | 0.93 | 0.91 | 0.57 | 0.73 | 0.51 | 0.91 | 0.95 | 0.93 | 0.59 |
| SJRHB013758_X2 | 0.94 | 0.92 | 0.53 | 0.85 | 0.75 | 0.72 | 0.87 | 0.79 | 0.54 |
| SJRHB013759_X1 | 0.97 | 0.93 | 0.51 | 0.81 | 0.62 | 0.72 | 0.97 | 0.96 | 0.55 |
| SJRHB015721_X1 | 0.92 | 0.91 | 0.57 | 0.79 | 0.79 | 0.69 | 0.95 | 0.93 | 0.80 |

Table S7: Summary of the predictive performance (Partial Least Squares) on each PDOXs cell line for training set, Leave-One-Out Cross-Validation (LOOCV) and test set.

| PDOXs cell line | Training Set | | | LOOCV | | | Test set | | |
|---|---|---|---|---|---|---|---|---|---|
| | R (Spearman) | $R^2$ | RMSE | R (Spearman) | $R^2$ | RMSE | R (Spearman) | $R^2$ | RMSE |
| SJEWS001321_X1 | 0.77 | 0.63 | 0.61 | 0.35 | 0.22 | 0.99 | 0.84 | 0.49 | 0.85 |
| SJMRT015723_X1 | 0.78 | 0.60 | 0.84 | 0.69 | 0.46 | 0.97 | 0.74 | 0.68 | 0.80 |
| SJNBL012407_X1 | 0.44 | 0.23 | 1.27 | 0.50 | 0.18 | 1.32 | 0.33 | 0.05 | 1.42 |
| SJNBL013762_X1 | 0.67 | 0.44 | 1.14 | 0.34 | 0.08 | 1.37 | 0.67 | 0.51 | 1.08 |
| SJNBL046_X | 0.62 | 0.37 | 1.11 | 0.69 | 0.55 | 0.97 | 0.61 | 0.27 | 1.15 |
| SJNBL108_X | 0.75 | 0.57 | 0.88 | 0.60 | 0.39 | 1.05 | 0.61 | 0.22 | 1.17 |
| SJOS001105_X1 | 0.68 | 0.48 | 0.69 | 0.62 | 0.58 | 0.69 | 0.75 | 0.51 | 1.07 |
| SJOS001107_X1 | 0.58 | 0.36 | 1.14 | 0.62 | 0.37 | 1.16 | 0.70 | 0.33 | 1.04 |
| SJOS001107_X2 | 0.64 | 0.54 | 0.72 | 0.45 | 0.45 | 0.82 | 0.39 | 0.58 | 0.97 |
| SJOS001107_X3 | 0.53 | 0.32 | 0.89 | 0.47 | 0.21 | 1.01 | 0.45 | 0.28 | 1.02 |
| SJOS001108_X1 | 0.40 | 0.22 | 1.12 | 0.43 | 0.42 | 1.12 | 0.70 | 0.45 | 1.36 |
| SJOS001112_X1 | 0.73 | 0.62 | 0.82 | 0.60 | 0.44 | 0.97 | 0.40 | 0.52 | 1.05 |
| SJOS010929_X1 | 0.48 | 0.29 | 1.29 | 0.58 | 0.36 | 1.26 | 0.66 | 0.42 | 1.50 |
| SJOS013769_X1 | 0.59 | 0.47 | 0.93 | 0.39 | 0.13 | 1.34 | 0.76 | 0.44 | 0.98 |
| SJRHB_TB-15-5700 | 0.86 | 0.76 | 0.80 | 0.52 | 0.29 | 1.59 | 0.79 | 0.66 | 0.93 |
| SJRHB000026_X1 | 0.70 | 0.54 | 1.08 | 0.64 | 0.38 | 1.21 | 0.71 | 0.52 | 1.11 |
| SJRHB000026_X2 | 0.62 | 0.34 | 1.00 | 0.69 | 0.44 | 1.00 | 0.72 | 0.58 | 0.97 |
| SJRHB010463_X16 | 0.49 | 0.25 | 1.11 | 0.60 | 0.47 | 1.02 | 0.38 | 0.23 | 1.43 |
| SJRHB010468_X1 | 0.54 | 0.47 | 0.78 | 0.52 | 0.41 | 0.90 | 0.66 | 0.42 | 1.06 |
| SJRHB010927_X1 | 0.75 | 0.55 | 0.99 | 0.63 | 0.51 | 1.12 | 0.77 | 0.53 | 1.48 |
| SJRHB011_X | 0.57 | 0.33 | 1.11 | 0.56 | 0.34 | 1.01 | 0.61 | 0.32 | 0.89 |
| SJRHB011_Y | 0.64 | 0.40 | 1.36 | 0.45 | 0.17 | 1.71 | 0.64 | 0.32 | 1.63 |
| SJRHB012_X | 0.44 | 0.23 | 1.30 | 0.63 | 0.36 | 1.17 | 0.41 | 0.22 | 1.76 |
| SJRHB012_Y | 0.75 | 0.59 | 1.08 | 0.42 | 0.15 | 1.45 | 0.66 | 0.56 | 1.16 |

| | | | | | | | | | |
|---|---|---|---|---|---|---|---|---|---|
| SJRHB012_Z | 0.49 | 0.18 | 1.39 | 0.45 | 0.17 | 1.51 | 0.41 | 0.19 | 1.29 |
| SJRHB012405_X1 | 0.63 | 0.36 | 0.94 | 0.61 | 0.35 | 0.95 | 0.67 | 0.45 | 1.12 |
| SJRHB013_X | 0.55 | 0.26 | 1.16 | 0.47 | 0.23 | 1.17 | 0.60 | 0.24 | 1.19 |
| SJRHB013758_X2 | 0.76 | 0.56 | 1.01 | 0.50 | 0.24 | 1.26 | 0.64 | 0.31 | 0.99 |
| SJRHB013759_X1 | 0.56 | 0.32 | 1.11 | 0.61 | 0.31 | 0.98 | 0.61 | 0.35 | 1.13 |
| SJRHB015721_X1 | 0.58 | 0.48 | 0.97 | 0.62 | 0.58 | 0.91 | 0.67 | 0.46 | 1.32 |
